# STAIG: Spatial Transcriptomics Analysis via Image-Aided Graph Contrastive Learning for Domain Exploration and Alignment-Free Integration

**DOI:** 10.1101/2023.12.18.572279

**Authors:** Yitao Yang, Yang Cui, Xin Zeng, Yubo Zhang, Martin Loza, Sung-Joon Park, Kenta Nakai

## Abstract

Spatial transcriptomics is an essential application for investigating cellular structures and interactions and requires multimodal information to precisely study spatial domains. Here, we propose STAIG, a novel deep-learning model that integrates gene expression, spatial coordinates, and histological images using graph-contrastive learning coupled with high-performance feature extraction. STAIG can integrate tissue slices without prealignment and remove batch effects. Moreover, it was designed to accept data acquired from various platforms, with or without histological images. By performing extensive benchmarks, we demonstrated the capability of STAIG to recognize spatial regions with high precision and uncover new insights into tumor microenvironments, highlighting its promising potential in deciphering spatial biological intricates.

## Introduction

Biological tissues are intricate networks of various cell types that perform essential functions through their unique spatial configurations. Recent spatial transcriptomics (ST) techniques, such as 10x Visium^1^, Slide-seq^2^, Stereo-seq^3^, and STARmap^4^, have greatly enhanced our ability to map genetic data within these configurations, providing deeper insights into the genetic organization of tissues in health and disease, and advancing our understanding of molecular and physiological intricacies.

ST heavily relies on the identification of spatial domains with uniform gene expression and histology. Currently, two main identification approaches exist: nonspatial and spatial clustering approaches^5^. Non-spatial clustering methods such as K-means^6^, Louvain^7^, and Seurat^8^ derive clustering results based solely on gene expression, often resulting in disjointed clusters that poorly reflect the actual spatial patterns of tissues. Emerging spatial clustering methods such as BayesSpace^9^ employ Bayesian models to integrate spatial correlations, whereas SEDR^10^ maps gene data using a deep autoencoder combined with spatial information through a variational graph autoencoder. STAGATE^11^ utilizes an attention mechanism in a graph autoencoder for adaptive learning of spatial and gene expression data. GraphST^12^ leverages graph neural networks with contrastive learning for spot representation. However, these methods do not utilize histological images such as Hematoxylin and Eosin (H&E)-stained images, which often result in unclear cluster boundaries and expert annotation misalignment.

Considering these limitations, new methods have been developed to integrate imaging data. stLearn^13^ offers a clustering framework that uses image-derived histological features for the normalization of gene expression. SpaGCN^14^ merges gene expression with spatial and histological pixel data in a weighted graph for graph convolutional network (GCN) analysis. DeepST^15^ extracts image features using a neural network to enhance the gene matrices. ConST^16^ uses the MAE^17^ visual model to extract histological features and merges them with gene data for domain identification using graph contrastive learning. However, the limited availability of histological images constrains the pre-training of deep models, diminishing their broad applicability and the quality of feature extraction. Furthermore, the extracted features, which primarily focus on color, are often distorted by inconsistent staining, thus limiting their analytical value.

In addition, recent graph-based approaches for ST data are flawed, as artificially structured graphs can lead to incorrect neighbor connections. This inaccuracy can disrupt the GCNs, compromising the precision of the results. Furthermore, methods integrating multiple tissue slices, the so-called batch integration, require manual alignment of slice coordinates and often overlook the subtle disparities between tissue slices, which still presents challenges for effective integration.

In this study, we propose STAIG (Spatial Transcriptomics Analysis via Image-Aided Graph Contrastive Learning), a deep learning framework based on the alignment-free integration of gene expression, spatial data, and histological images, to ensure refined spatial domain analyses. STAIG extracts features from H&E-stained images using a self-supervised model and builds a spatial graph with these features. The graph is further processed via contrastive learning using a graph neural network (GNN) that generates informative embeddings. We evaluated the performance of STAIG using diverse datasets and observed its promising capability for spatial domain identification. We demonstrated that STAIG revealed the spatial and genetic details of tumor microenvironments, advancing our understanding of complex biological systems.

## Results

### Overview of STAIG

To reduce noise and uneven staining, STAIG first segments histological images into patches that align with the spatial locations of data spots, and subsequently refines these images with a band-pass filter. Image embeddings were extracted using the Bootstrap Your Own Latent (BYOL)^18^ self-supervised model, and an adjacency matrix was simultaneously constructed based on the spatial distances between the spots (Fig. 1a). To integrate the data from different tissue slices, the image embeddings for multiple tissue slices were stacked vertically (Fig. 1b), and the adjacency matrices were merged using a diagonal placement method (Fig. 1c).

**Fig. 1.**
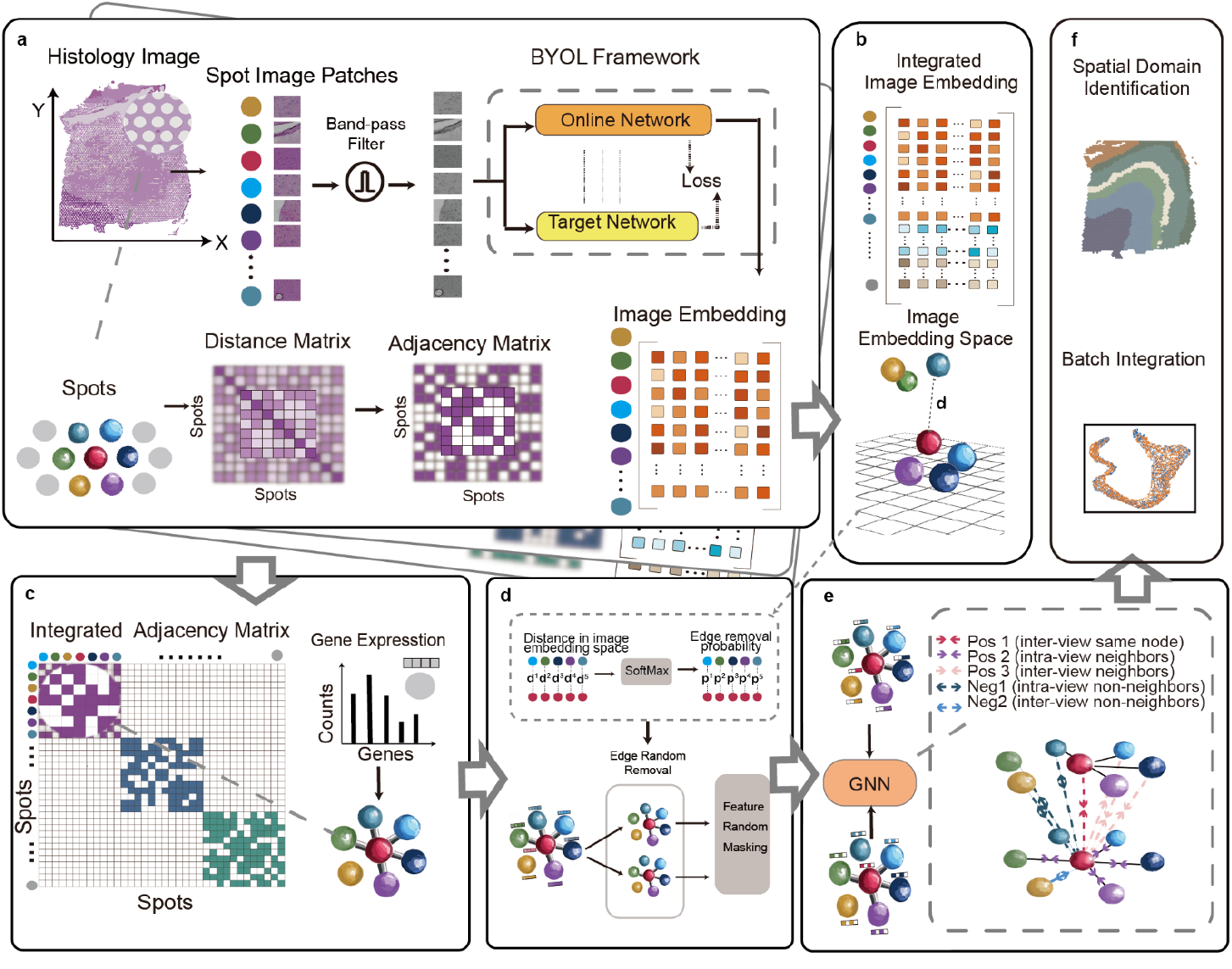
Overview of the STAIG framework. (**a**) STAIG begins with spatial transcriptomic (ST) data. Each slice includes spots with spatial coordinates, gene data, and optional Hematoxylin and Eosin (HE) stained images. Image patches at these spots undergo noise reduction, including bandpass filtering, before being processed for image embeddings via the BYOL framework. Parallelly, an adjacency matrix is created using the spatial data of the spots. (**b**) For multiple slices, the image embeddings from each slice are vertically merged, forming an image embedding space where spots are distributed, with dotted lines indicating their Euclidean distances. (**c**) Adjacency matrices from each section are combined diagonally to form an integrated adjacency matrix. This matrix is then used to construct a graph, with gene expression data represented as node information. (**d**) For spots connected by edges, distances are calculated in the image embedding space. These distances are then transformed into probabilities of random edge removal using a SoftMax function. The original graph undergoes two rounds of edge random removal based on these probabilities, creating two augmented views. Subsequently, features of nodes in these graph views are randomly masked. (**e**) A graph neural network (GNN) processes these augmented views, concentrating on node-level differences. This step is encapsulated within a dotted box, emphasizing the neighbor contrastive strategy. (**f**) The derived embeddings from the GNN are then utilized for spatial domain identification and integration.

Next, an original graph was built from the integrated adjacency matrix, where nodes represent gene expression and edges reflect adjacency (Fig. 1c). To generate two independent augmented views from this graph, STAIG employs a parallel and independent augmentation process (Fig. 1d). For each augmented view, the process involved randomly removes edges from the original graph, guided by an image-driven probability (or a fixed probability in the absence of images). This is followed by the random masking of gene features, entailing the zeroing of a subset of gene values. More importantly, the probability of edge removal is estimated based on the Euclidean distance between nodes in the image feature space, thereby introducing image-informed augmentation into the graph.

The augmented views are processed using a shared GNN guided by a neighboring contrastive objective. This approach aims to closely align nodes and their adjacent neighbors in two graph views while distancing non-neighboring nodes (Fig. 1e). Additionally, when images are available, the Debiased Strategy (DS)^19^ leverages image embeddings as prior knowledge to ensure nodes that are closer based on image features remain proximate (details in the Method section). Finally, the trained GNN produced embeddings to identify the spatial regions and minimize batch effects across consecutive tissue slices (Fig. 1f).

### Performance in the identification of brain regions

To assess the performance of STAIG in the identification of tissue regions, we prepared 10x Visium human and mouse brain ST datasets: 12 human dorsolateral prefrontal cortex (DLPFC)^20^ slices that were manually annotated into cortical layers L1‒ L6 and white matter (WM), and mouse anterior and posterior brain sections. We benchmarked STAIG with existing methods such as Seurat, GraphST, DeepST, STAGATE, SpaGCN, SEDR, ConST, and stLearn. To quantify the performance, the Adjusted Rand Index (ARI)^21^ for manually annotated datasets and the Silhouette Coefficient (SC)^22^ for others were employed (refer to ‘Baseline methods and evaluation metrics’ in the Supplementary Notes).

Overall, for the human brain datasets, STAIG achieved the highest median ARI (0.69 across all slices) (Fig. 2a, Supplementary Fig. S1 and S2); in particular, the ARI reached 0.84 in slice #151672 (Fig. 2b). Regarding slice #151673 used in previous studies^23–25^, STAIG achieved the highest ARI of 0.68, not only distinctly separating tissue layers L1‒L6 and WM in uniform manifold approximation and projection (UMAP) visualizations but also closely matched manual labels and accurately recognized layer proportions (Fig. 2c). Conversely, the results demonstrate that the existing methods provide relatively lower ARIs with misclassified spots and inaccuracies in layer proportions (Fig. 2d-e): stLearn yielded a missing layer and misclassifying spots; GraphST achieved an ARI of 0.64 but had discrepancies in the positioning of layers 4 and 5; other methods recorded ARIs between 0.46 and 0.54 owing to inaccuracies in layer proportions.

**Fig. 2.**
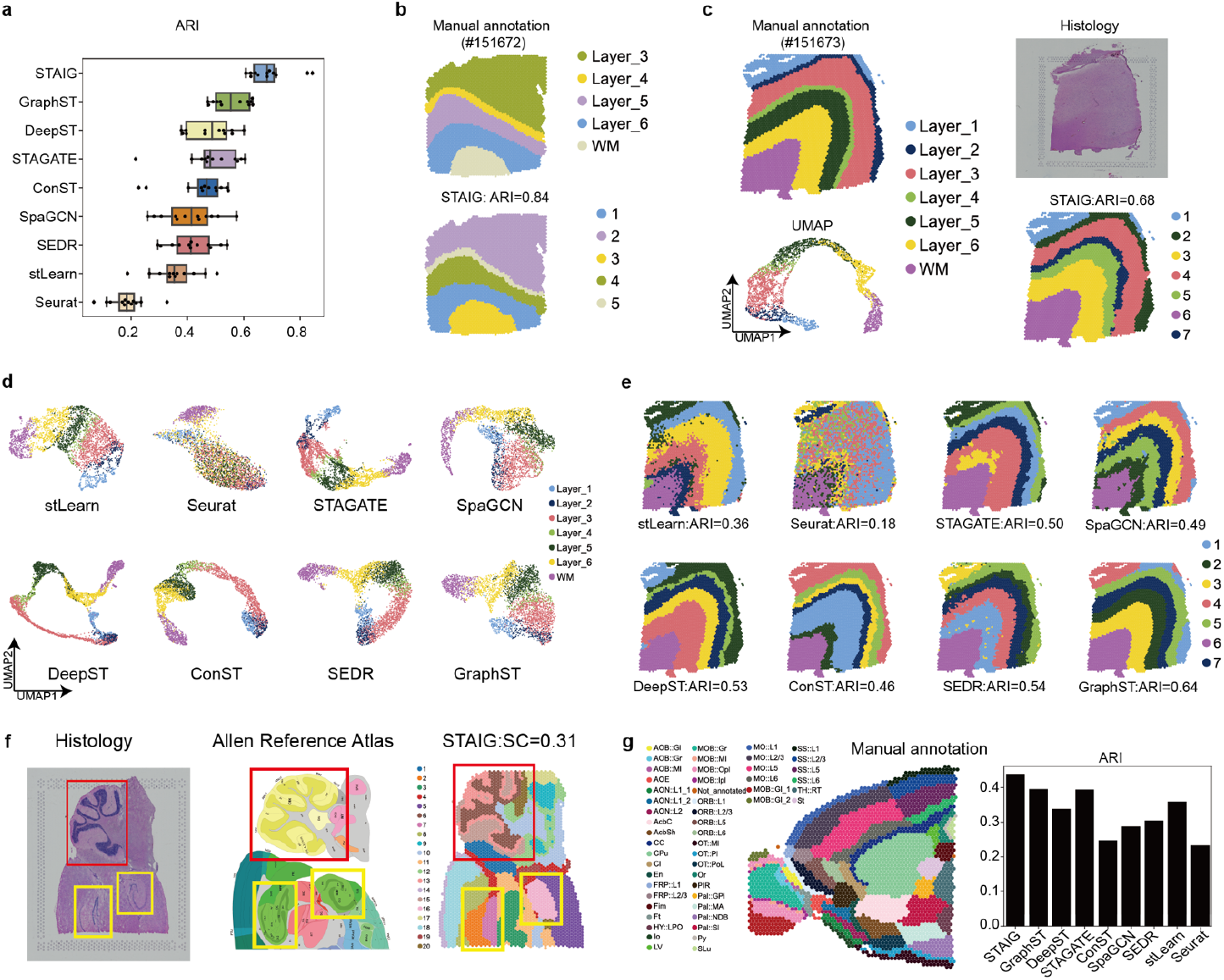
Enhanced spatial region identification in Human Dorsolateral Prefrontal Cortex (DLPFC) and mouse brain tissues by STAIG. **(a)** Boxplots of adjusted rand index (ARI) of the nine methods applied to all 12 DLPFC slices. In the boxplot, the center line denotes the median, box limits denote the upper and lower quartiles, black dots denote individual slices, and whiskers denote the 1.5× interquartile range. **(b)** Manual annotations and clustering results with ARI by STAIG on DLPFC slice #151672. **(c)** Manual annotations, H&E stained image, clustering result with ARI and UMAP visualization by STAIG on DLPFC slice #151673. **(d)** UMAP visualizations by baseline methods (stLearn, Seurat, STAGATE, SpaGCN, DeepST, conST, SEDR, GraphST) on DLPFC slice #151673. **(e)** Clustering results with ARI by baseline methods on DLPFC slice #151673. Manual annotations and clustering results of the other DLPFC slices are shown in Supplementary Fig. S1 and S2. **(f)** H&E stained image, anatomical annotations from the Allen Reference Atlas, and clustering results with Silhouette Coefficient (SC) by STAIG on mouse brain posterior tissue. The red box denotes the cerebellar cortex, and the yellow box denotes the hippocampus area with Cornu Ammonis (CA) and Dentate Gyrus (DG). Clustering results from baseline methods are shown In Supplementary Fig. S3. **(g)** Manual annotations from Long et al. and bar charts of ARI by STAIG and baseline methods on mouse anterior tissue. The y-axis represents ARI. Clustering results from baseline methods are shown in Supplementary Fig. S4.

In the mouse posterior tissue dataset (Fig. 2f and Supplementary Fig. S3), STAIG successfully identified the cerebellar cortex and hippocampus region, further distinguishing the Cornu Ammonis (CA) and dentate gyrus sections. This was consistent with the Allen Mouse Brain Atlas annotations^26^. Although the overall accuracy decreased in the absence of manual labels, STAIG achieved the highest SC of 0.31. In the mouse anterior tissue dataset (Fig. 2g and Supplementary Fig. S4), STAIG accurately demarcated areas, including the olfactory bulb and dorsal pallium, yielding the highest ARI of 0.44 when we used manual labels from Long et al.^12^ as a reference.

### Efficacy of image feature extraction

To investigate the impact of the usage of image features, we applied the k-nearest neighbors algorithm (KNN)^27^ to the image features from STAIG and those from the comparison algorithms, that is, stLearn, DeepST, and ConST. For instance, in the analysis of slice #151507 (Fig. 3a), we found that the image features from stLearn were heavily influenced by stain intensity and led to mismatches with the actual layer annotations. Despite utilizing deep learning models, DeepST and ConST failed to capture the intricate texture characteristics of brain tissues.

**Fig. 3.**
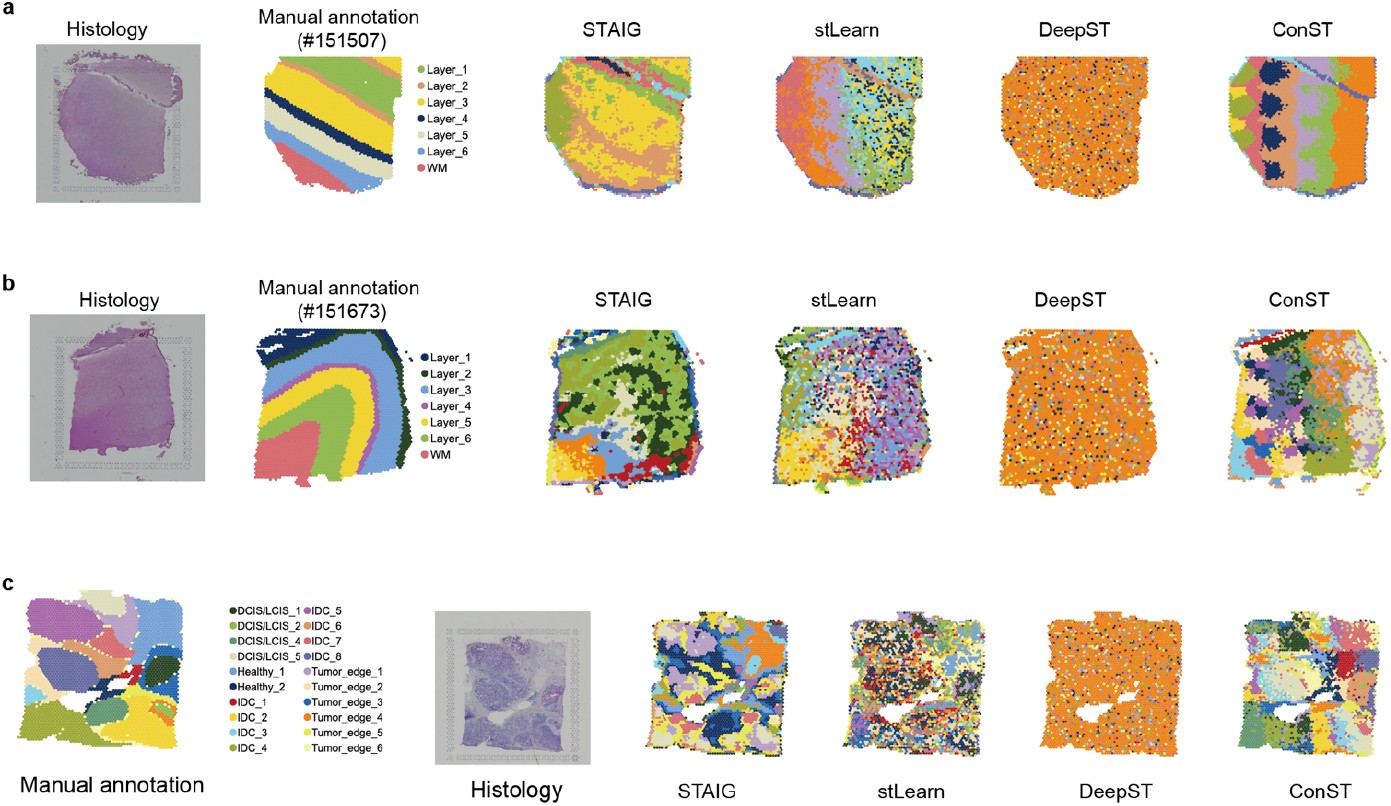
Improving spatial transcriptomic analysis through advanced image feature extraction in STAIG. **(a)** H&E stained image, manual annotations, and KNN clustering results based purely on image features by STAIG and three other image-based methods (stLearn, DeepST, ConST) on DLPFC slice #151507. **(b)** H&E stained image, manual annotations and KNN clustering results based purely on image features by STAIG and three other image-based methods on DLPFC slice #151673. **(c)** H&E stained image, visual interpretation-based manual annotations, and KNN clustering results based purely on image features by STAIG and three other image-based methods on human breast cancer dataset.

Remarkably, STAIG exhibited a high degree of conformity with the texture distribution in the manually annotated layers, with only minor deviations attributable to staining variations (Fig. 3a). This precision was particularly evident in the clear demarcation between the WM and Layer 6, as well as in the accurate distribution of Layer 3 on either side of Layer 1. Furthermore, the performance of STAIG image features on slice #151673 was equally impressive (Fig. 3b). Interestingly, the comparative methods for this slice generally succeeded in differentiating only between the WM and non-WM layers, whereas the clustering of STAIG within the non-WM layers aligned closely with manual annotations, particularly for layer 4.

We further investigated the effectiveness of the image features using a human breast cancer H&E-stained image (Fig. 3c); a pathologist manually annotated regions solely based on visual interpretation^10^. The results indicated that the image features from stLearn mixed tumor and normal regions while those from ConST displayed a significant deviation from manual annotations. DeepST failed to capture image features. Conversely, STAIG clustering precisely identified the tumor regions and maintained spatial region coherence in the clustering results, suggesting advanced image feature extraction in STAIG.

### Identifying spatial domains without histological image data

Because ST experiments lacking image information are not unusual, we assessed the predictive performance with datasets where only gene expression and spatial information were available. Initially, we prepared a mouse coronal olfactory bulb dataset of Stereo-seq^3^ annotated using DAPI-stained images^10^ (Fig. 4a): the olfactory nerve layer (ONL), glomerular layer (GL), external plexiform layer (EPL), mitral cell layer (MCL), internal plexiform layer (IPL), granule cell layer (GCL), and rostral migratory stream (RMS).

**Fig. 4.**
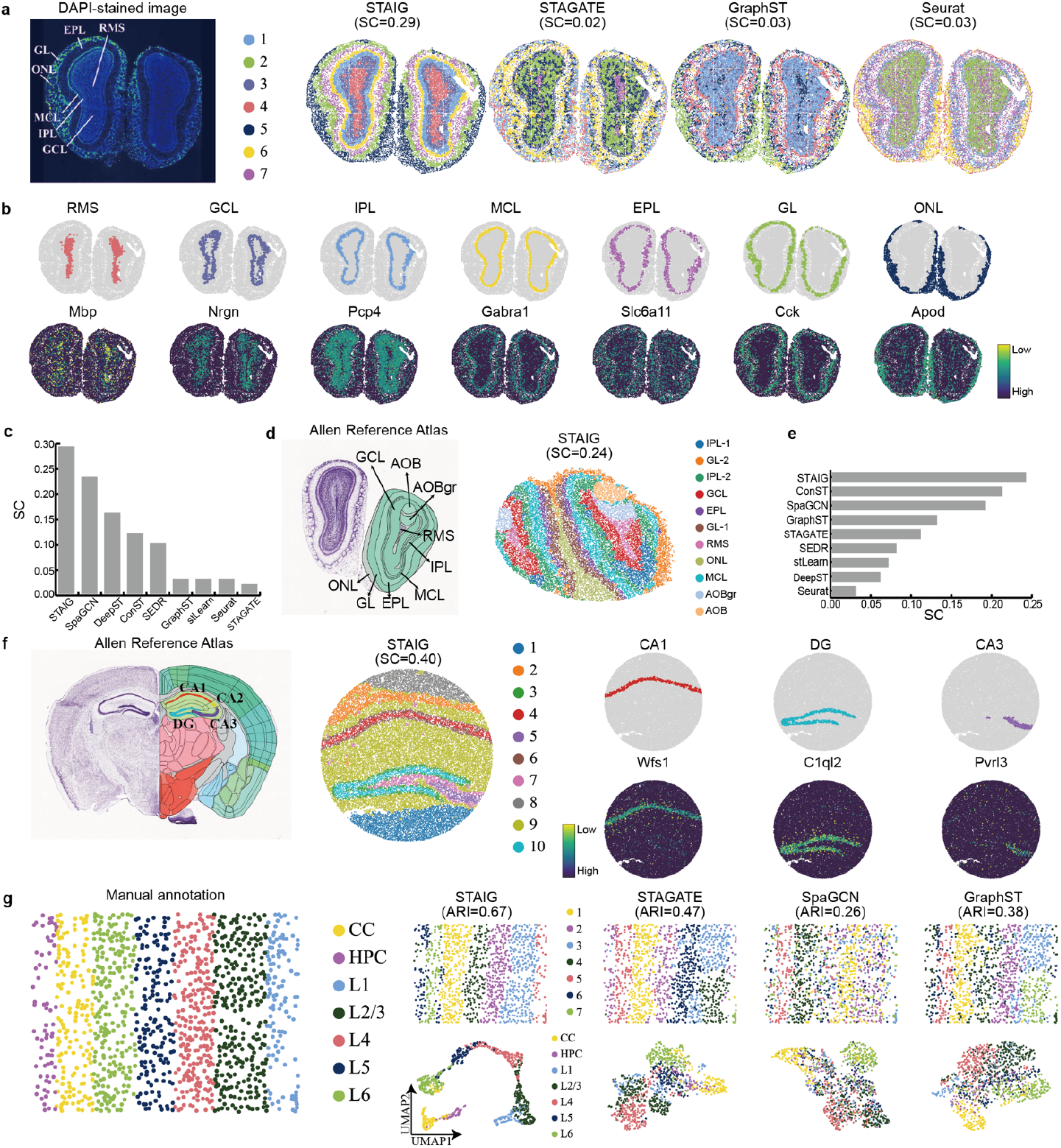
Robust performance of STAIG in spatial transcriptomics lacking image data. (**a**) DAPI-stained image with annotated laminar organization and clustering results with SC by STAIG, STAGATE, GraphST, and Seurat on Stereo-seq mouse olfactory bulb tissue. Clustering results from other baseline methods are shown in Supplementary Fig. S5. (**b**) Visualization of spatial domains identified by STAIG and the corresponding marker gene expressions on the Stereo-seq mouse olfactory bulb tissue. (**c**) Bar chart of SC by STAIG and baseline methods on Stereo-seq mouse olfactory bulb tissue. The y-axis represents SC. (**d**) Annotations from the Allen Reference Atlas and clustering results with SC by STAIG on Slide-seqV2 mouse olfactory bulb tissue. Clustering results from baseline methods are shown in Supplementary Fig. S6. (**e**) Bar chart of SC by STAIG and baseline methods on Slide-seqV2 mouse olfactory bulb tissue. The x-axis represents SC. (**f**) Annotations from the Allen Reference Atlas, clustering results with SC, and visualization of CA1, CA3, and DG domains identified by STAIG and the corresponding marker gene expressions on Slide-seqV2 mouse hippocampus tissue. Clustering results from baseline methods are shown in Supplementary Fig. S7. (**g**) Manual annotations and clustering results with ARI by STAIG, STAGATE, GraphST, and Seurat on STARmap mouse visual cortex dataset. Clustering results from other baseline methods are shown in Supplementary Fig. S8.

All algorithms provided the overall trends of spatial stratification for the dataset, whereas disparities in the annotations were observed (Fig. 4a, Supplementary Fig. S5). For example, Seurat and stLearn fail to distinguish between layers, and STAGATE and GraphST mix the IPL and GCL. DeepST yielded distinct stratifications but failed to discern the GL and IPL layers. Remarkably, STAIG successfully delineated all layers, including thinner layers such as the MCL and IPL. As shown in Fig. 4b, the expression of layer-specific marker genes^28–33^ supported the high-resolution predictions of STAIG, outperforming the existing algorithms (Fig. 4c).

Next, we tested the algorithms using the mouse olfactory bulb tissue dataset Slide-seqV2^34^ (Fig. 4d and Supplementary Fig. S6a). All algorithms, except STAIG, GraphST, and STAGATE, struggled to discern the stratification of the RMS, accessory olfactory bulb (AOB), and granular layer of the accessory olfactory bulb (AOBgr). In contrast, STAIG displayed a higher performance in interlayer clustering (Fig. 4e), and the results of STAIG matched well with the expression of layer-specific marker genes^35,36^ (Supplementary Fig. S6b).

Using the mouse hippocampus dataset^34^ from Slide-seqV2 (Fig. 4f and Supplementary Fig. S7), were further assessed (Fig. 4f and Supplementary Fig. S7). Notably, several methods, such as Seurat, stLearn, SpaGCN, and SEDR, display unclear clustering results that lack spatial cohesion. However, STAIG, GraphST, STAGATE, and DeepST delineated spatially consistent clusters for the major anatomical regions, particularly the CA, which was accurately categorized into CA1 and CA3 sections. Again, STAIG had the highest SC of 0.40, which was validated by the expression of marker genes^37–39^.

Lastly, using the mouse visual cortex dataset of STARmap dataset^4^ with inherent annotations, we found that STAIG outperformed all other methods, achieving the highest ARI of 0.67, which markedly surpassed the next-best method, STAGATE (Fig. 4g and Supplementary Fig. S8). Moreover, the UMAP plots highlight the unique ability of STAIG to completely separate all neocortical layer clusters.

### Rebuilding consecutive slices without prior spatial alignment

To test the performance of the vertical slice integration, we used 12 sections of the DLPFC acquired from three individuals. Overall, the existing methods were less effective, particularly for slices #151508 and #151509 spaced 300 μm apart (Supplementary Fig. S9 and S10). DeepST failed to separate the clusters from layers 3‒5, and GraphST insufficiently separated the layers in the UMAP representation. In contrast, the STAIG successfully integrated these DLPFC slices at various heights (Fig. 5a-b). It achieved complete UMAP overlap, reflected in an ARI of 0.60 for slices merely 10 μm apart (#151675 and #151676), and 0.58 for those 300 μm apart.

**Fig. 5.**
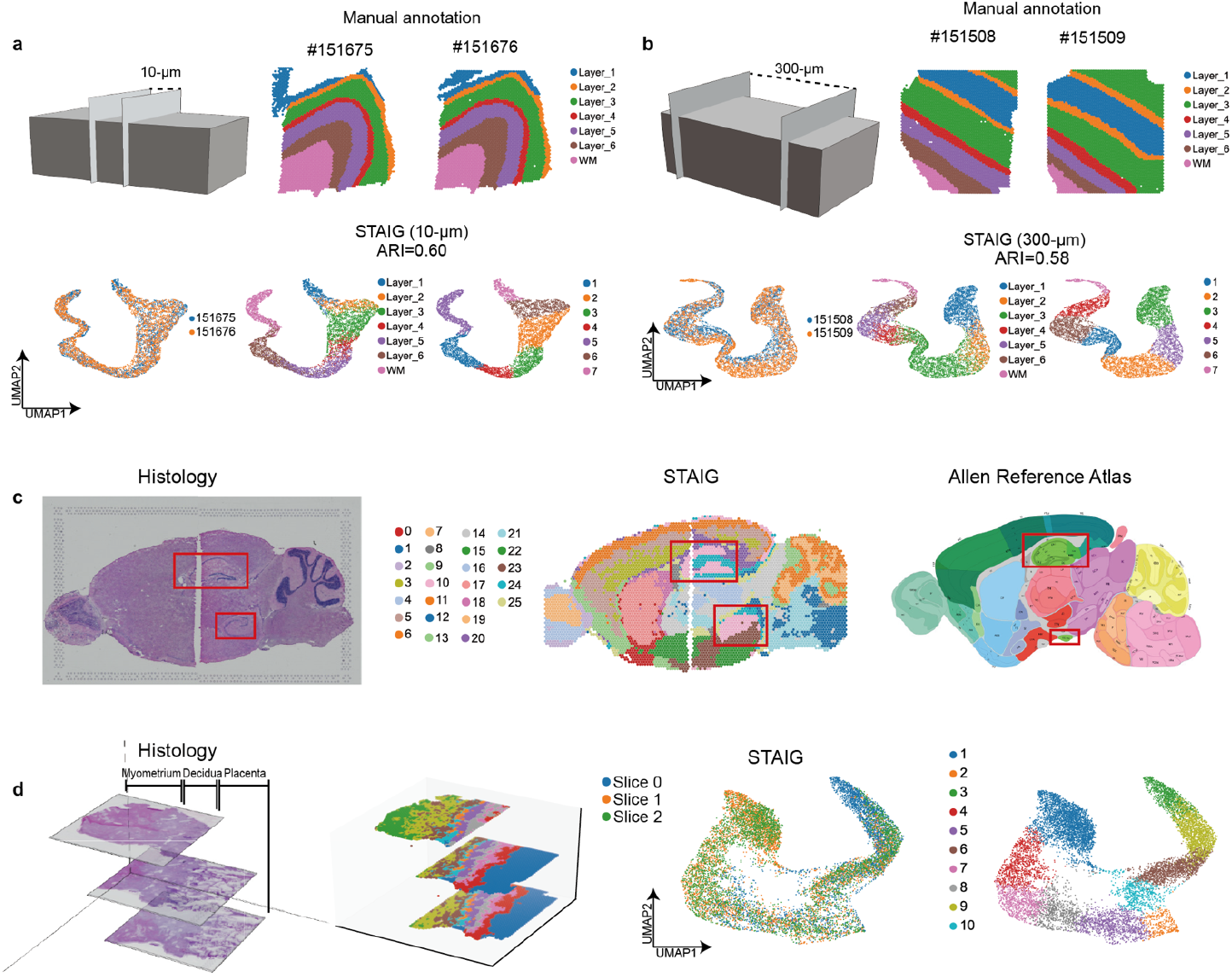
Integration Capabilities of STAIG. **(a)** Vertical integration of STAIG at 10 μm (DLPFC slice #151675 and #151676). Includes a simplified slice sampling diagram, manual annotations, integrated UMAP, and ARI. Baseline methods results are detailed in Supplementary Figure S9. **(b)** Vertical integration of STAIG at 300 μm (DLPFC slice #151508 and #151509). Includes a simplified slice sampling diagram, manual annotations, integrated UMAP, and ARI. Baseline methods results are detailed in Supplementary Fig. S10. (**c**) Horizontal integration of STAIG. Includes H&E stained images, annotations from the Allen Reference Atlas and integration results by STAIG on mouse anterior and posterior brain tissue slices. The red box highlights the hippocampus zones in both slices. Baseline methods results are shown in Supplementary Fig. S11. (**d**) Partial overlap integration of STAIG. H&E stained images and integration results with UMAP visualization by STAIG on three partially overlapping slices from the human placental bed dataset.

Anterior and posterior mouse brain slices were used to test methods using horizontal slice integration. Unlike the methods SpaGCN and STAGATE, which require pre-alignment of slice edges, STAIG without such prior spatial alignment precisely identified key brain structures, such as the cerebral cortex layers, cerebellum, and hippocampus, which aligned well with the annotation of the Allen Brain Atlas (Fig. 5c) and showed accuracy comparable to the existing methods (Supplementary Fig. S11).

To further investigate the performance of partial overlap integration, we prepared the human placental bed dataset, in which three consecutive overlapping slices from a single donor, including the Myometrium, Decidua, and Placenta, were segmented into 10 clusters^40^. Remarkably, the STAIG seamlessly integrated the slices and eliminated batch effects, as shown in the UMAP plots (Fig. 5d). In particular, slice 0 distinctively showed domain 3, which is unique to the internal tissue of the myometrium. This distribution was exclusive to slice 0 and absent in slices 1 and 2. These results demonstrate that STAIG enables not only the assimilation of overlapping regions but also the maintenance of the distinct characteristics of unique spatial regions.

### Specifying the tumor microenvironment from human breast cancer ST

In the analysis of the human breast cancer dataset, we found that the STAIG results closely matched the manual annotations and achieved the highest ARI of 0.64 (Supplementary Fig. S12). Notably, STAIG proposed a slightly different, yet more refined spatial stratification. In particular, for the region ‘Healthy_1’ in the manual annotation (Fig. 6a), STAIG dissected it into subclusters Clusters 3 and 4 (Fig. 6b). We noticed that the differentially expressed genes (DEGs) in Cluster 3 compared with other clusters (>0.25 log2FoldChange, denotes as log2FC) (Fig. 6c) were significantly involved in biological processes, such as extracellular matrix organization, wound healing, and collagen fibril organization (Fig. 6d).

**Fig. 6.**
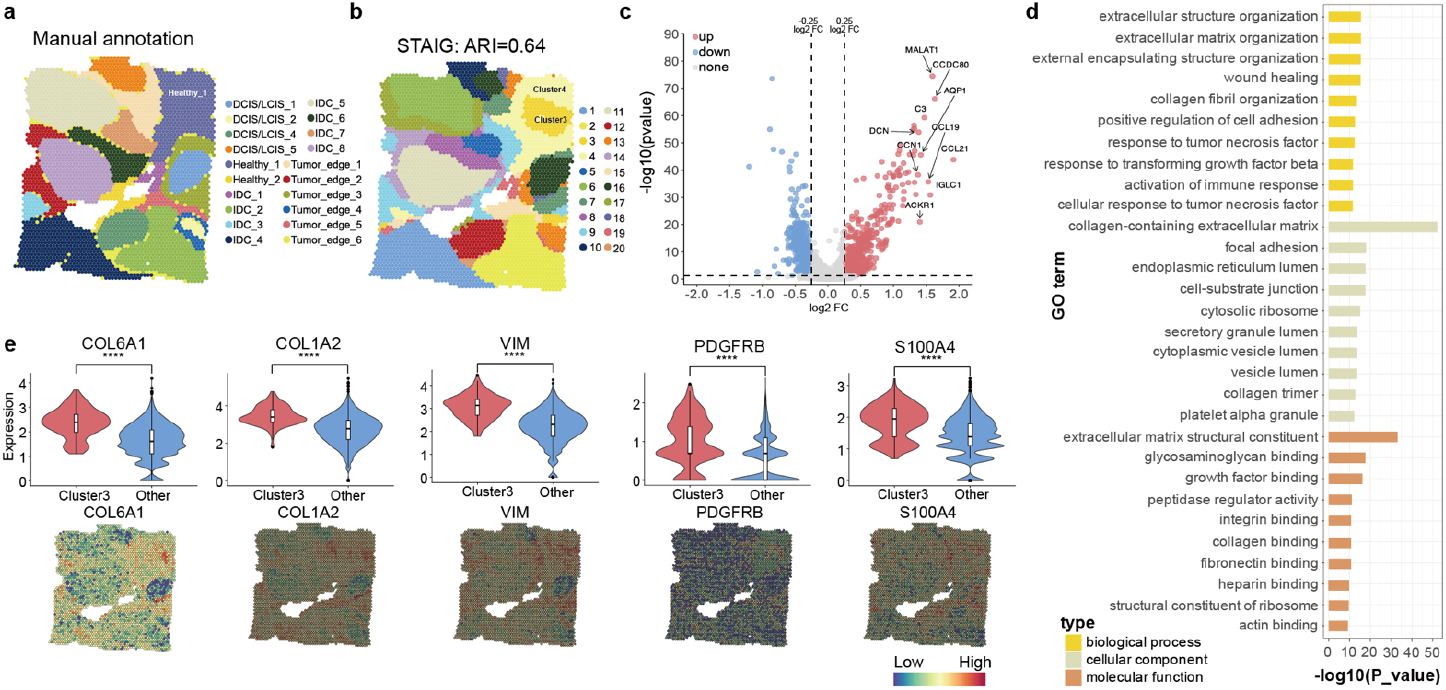
Advanced spatial analysis by STAIG reveals Cancer-associated fibroblasts (CAF)-rich clusters in human Breast Cancer ST Data. (**a**) Manual annotation of human breast cancer dataset based on the HE-stain image. (**b**) Clustering results with ARI by STAIG on human breast cancer dataset. Clustering results from other baseline methods are shown in Supplementary Fig. S12. (**c**) Differential Gene Expression (DGE) analysis of Cluster 3 versus other clusters. Each point represents a gene, the vertical axis represents the -log10 of the p-value and the horizontal axis represents the log2FoldChange (log2 FC). The significance threshold is set at |log2FC| > 0.25. (**d**) Gene Ontology (GO) analysis for Cluster 3 versus other clusters. The vertical axis represents the GO terms, and the horizontal axis represents the -log10 of the p-value. (**e**) Violin plots and the visualization of expression of CAF marker genes (*COL*6*A*1, *COL*1*A*2, *VIM, PDGFRB, S*100*A*4) in Cluster 3 versus other clusters. The vertical axis represents gene expression levels. Each ‘violin’ represents the distribution of expression for a particular gene, with the width indicating frequency. The large box between pairs of violins shows the statistical comparison between Cluster 3 and other clusters. The presence of stars in this box denotes statistical significance, with the number of stars corresponding to the level of significance (e.g., *p < 0.05, **p < 0.01, ***p < 0.001, ****p < 0.0001).

Interestingly, DEGs included *DCN, CCL*19, and *CCL*21, which are known to function in cancer-associated fibroblasts (CAFs)^41,42^. Moreover, the CAF marker genes (e.g., *COL*6*A*1, *COL*1*A*2, *VIM, PDFGRB, S100A4*)^43–45^ were significantly upregulated in Cluster 3 (Fig. 6e) but not in Cluster 4 (Supplementary Fig. S13a and S13b).

Taken together, through the integration of multimodality by STAIG, we found that Cluster 3 shapes a tumor microenvironment densely populated by CAFs and revealed the molecular properties of CAF-rich areas.

### Identifying tumor-adjacent tissue junctions in zebrafish melanoma ST

To further characterize the tumor microenvironment, we investigated the A and B tissue slices of zebrafish melanoma (Fig. 7a). The previous study incorporated scRNA-seq data, thereby segmenting the interface clusters into muscle-like and tumor-like subclusters^46^. These clusters could not be specified using ST data analysis alone. Remarkably, STAIG identified the interfaces at the tumor-adjacent tissue junction using only ST data (Fig. 7b), which often failed in traditional ST methods (Supplementary Fig. S14, S15). In addition, the DEGs found at the interfaces included known genes such as *si*: *dkey* −153*m*14.1, *zgc*: 158463, *RPL*41, and *hspb*9 (Fig. 7c and Supplementary Fig. S16a) and were associated with mRNA metabolic processes (Fig. 7d and Supplementary Fig. S16b). These results highlight the active transcriptional and translational processes in these regions, thus supporting previous findings.

**Fig. 7.**
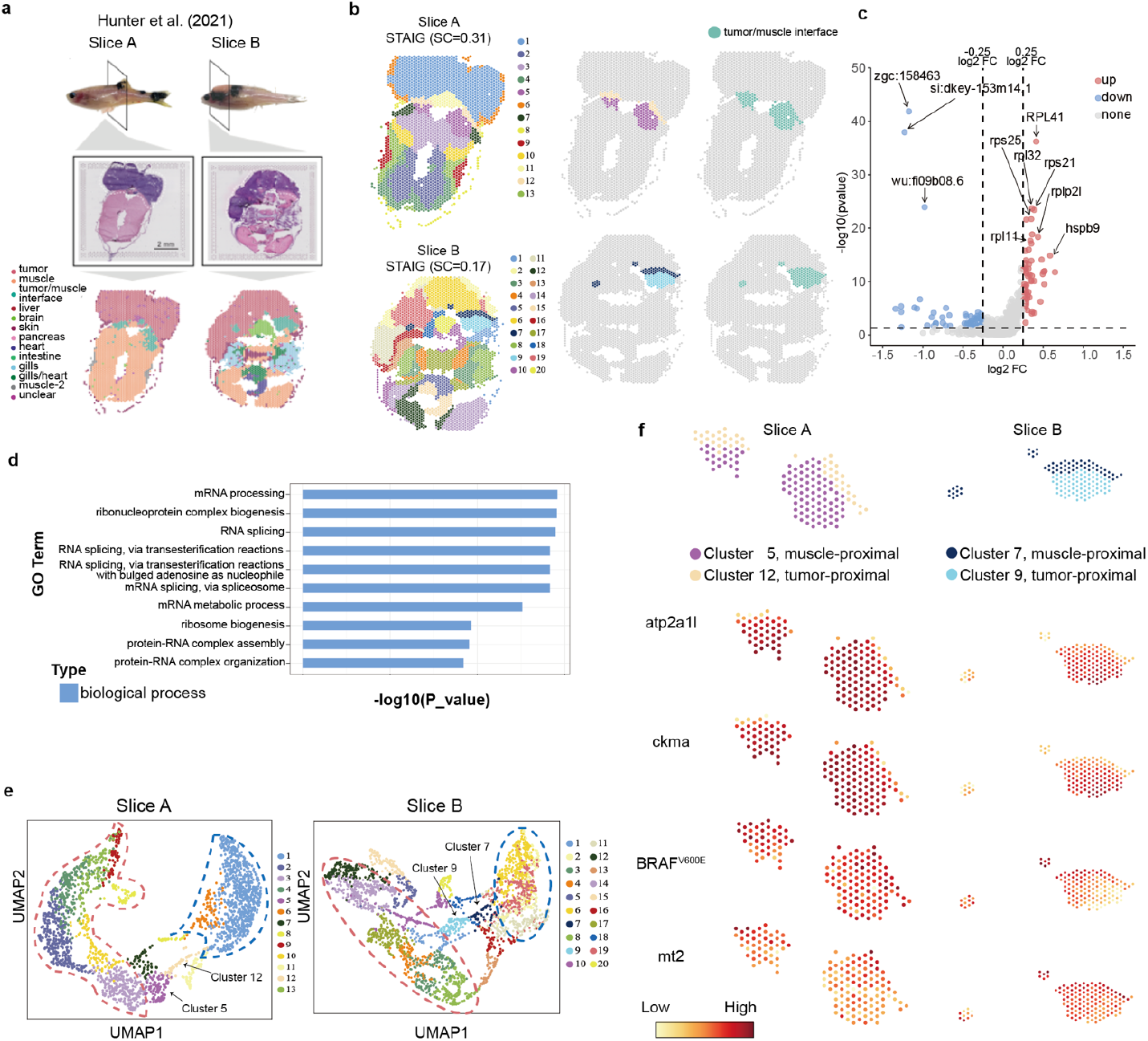
Enhanced resolution of tumor interface dynamics in zebrafish melanoma using STAIG analysis. (**a**) H&E stained image and annotation of zebrafish melanoma on slices A and B from Hunter et al. (2021). (**b**) Interface domains identified by STAIG on slices A and B with SC. Baseline methods results are shown in Supplementary Fig. S13 and S14. (**c**) DGE analysis of the interface domain versus other domains on slice A. (**d**) GO analysis for the identified interface domain versus other domains on slice A. (**e**) UMAP visualization of clustering results on slice A and B by STAIG. The red outline indicates clusters of normal tissue, while the blue outline denotes clusters of tumor tissue. (**f**) Fine-tuned subregions (muscle-proximal and tumor-proximal) of the interface identified by STAIG and the gene expression patterns of specific genes (*atp*2*a*1*l, ckma, BRAF*^*V*600*E*^, *mt*2).

Subsequently, STAIG identified Cluster 12 in slices A and 7 in slice B as being closer to the tumor area, whereas Cluster 5 in slices A and 9 in slice B were positioned nearer to the muscle tissue. As shown in the UMAP plots (Fig. 7e), although Clusters 12 and 5 in slice A were interconnected, Cluster 12 was aligned with tumor-associated clusters, and Cluster 5 had normal tissue clusters. A similar pattern was observed in Clusters 7 and 9 in slice B. Notably, the muscle-proximal subcluster showed high expression of muscle marker genes^47,48^, such as *atp*2*a*1*l* and *ckma* (Fig. 7f and Supplementary Fig. S16c-d), and the tumor-proximal subcluster exhibited high expression of tumor-associated genes^49,50^, for example, *BRAF*^*V*600*E*^ and *mt*2. These marker genes exhibited differential functional involvement (Supplementary Fig. S16e): the subcluster farther from the tumor showed an enrichment of muscle structure and metabolism, whereas those closer to the tumor were involved in protein folding and responding to temperature changes, which are indicative of increased cellular activity and stress responses typically found in tumor cells^51^.

In summary, we characterized the tumor borders of zebrafish melanoma and found that the cluster near tumors resembled a tumor-like interface, whereas that closer to normal tissue was akin to a muscle-like interface. These results highlight the necessity for ST annotation and dissection at a higher resolution to enhance our understanding of intricate cancer invasion.

## Discussion

In this study, we proposed STAIG, a novel deep learning model that efficiently integrates gene expression, spatial coordinates, and histological images using graph contrastive learning coupled with high-performance feature extraction. Unlike other methods, our approach first processes images using Gaussian blurring and bandpass filtering, which are essential for eliminating noise and enabling the model to focus on texture rather than color. Therefore, our model successfully extracted spatial features from an entire H&E-stained image without relying on additional image data. These features were processed by graph augmentation along with a unique contrastive loss function to focus on neighbor node consistency, which contributed to improving the performance of STAIG even in settings without image data.

Furthermore, the contrastive integration of augmented graphs focuses on local node adjustments to facilitate dimensionality reduction and maintain node-to-neighbor relationships across different slices. This strategy allows STAIG to process multiple slices together in an alignment-free manner by avoiding batch effects, as exemplified by the reconstruction of vertical, horizontal, and partially overlapping tissue slices. This integration is most effective when applied to multiple tissue slices from the same source, and requires further investigation to overcome this limitation.

By extensively benchmarking various ST datasets, we demonstrated the effectiveness of STAIG by utilizing advanced image feature extraction if histological images were available. Remarkably, in the human breast cancer and zebrafish melanoma datasets, STAIG not only identified spatial regions with high resolution compared to existing studies but also uncovered previously challenging-to-identify regions, offering unprecedented depth into tumor microenvironments. Additionally, STAIG showed exceptional capability in delineating tumor boundaries, outperforming others in identifying transitional zones with precision. This effectiveness is also linked to the contrast strategy of the STAIG, which captures vital local information.

In conclusion, we confirmed the impact of combining advanced machine learning with multimodal spatial information and the promising potential of STAIG in deciphering cellular architectures, which facilitates our understanding of spatial biological intricacies. In future work, because constructing a whole-graph representation is resource-intensive, especially in systems with limited GPU memory, optimizing matrix compression during graph construction and effectively applying mini-batch approaches to optimize GNNs should be further considered.

## Methods

### Data description

Publicly available ST datasets and histological images acquired from various platforms were downloaded (Supplementary Table S1). From the 10x Visium platform, the DLPFC dataset included 12 sections from three individuals, with each individual contributing four sections sampled at 10 μm and 300 μm intervals. The spot counts for these datasets varied between 3,498 and 4,789 per section, with the DLPFC dataset featuring annotations for its six layers and white matter. The human breast cancer dataset had 3,798 spots, and the mouse brain dataset comprised the anterior and posterior sections with 2,695 and 3,355 spots, respectively. For zebrafish melanoma, two tissue slices, A and B, with 2,179 and 2,677 spots, respectively, were analyzed. The human placental bed dataset included three consecutive slices from a single donor with 3,568, 3,855, and 4,186 spots per slice. For the integration experiments, the DLPFC, mouse brain, and human placental bed datasets were used.

The Stereo-seq dataset of the mouse olfactory bulb included 19,109 spots with a 14 μm resolution. Slide-seqV2 datasets featured a 10 μm resolution with the mouse hippocampus (18,765 spots from the central quarter radius) and the mouse olfactory bulb (19,285 spots). The STARmap dataset comprised 1,207 spots. These datasets lacked images, and image annotations were downloaded from the Allen Brain Atlas website (Supplementary Table S2).

### Data preprocessing

Raw gene expression counts were log-transformed and normalized using the SCANPY^52^ package. After normalization, the data were scaled to achieve a mean of zero and unit variance. For our experiments, we selected a subset of *F* highly variable genes (HVGs), where *F* was designated as 3,000.

### Basic graph construction

For each slice, we constructed a basic undirected neighborhood graph, *G* = (*V, E*), where *V* represents the set of *N* spots {*v*_1_, *v*_2_, … *v*_*N*_}, and *E* ⊆ *V* × *V* denotes the edges between these spots. The corresponding adjacency matrix, *A* ∈ {0,1}^*N*×*N*^, is defined such that *A*_*ij*_ = 1 indicates the presence of an edge (*v*_*i*_, *v*_*j*_) ∈ *E*. We determined the edges by computing a distance matrix *D* ∈ ℝ^*N*×*N*^, with *D*_*ij*_ representing the Euclidean distance between any two spots *v*_*i*_ and *v*_*j*_. KNN was employed to connect each spot with its top *k* closest spots, forming neighboring nodes. The gene expression matrix *X* ∈ ℝ^*N*×*F*^ was also established, where each row corresponds to the expression profile of HVGs for each spot.

### Histological image feature extraction

For the ST data with H&E-stained slices, the extraction of image features from each spot was critical. We started with Gaussian blurring of the entire H&E-stained image to reduce texture noise and staining biases. Next, 512 × 512 pixel patches centered on the spot coordinates were extracted. These *N* patches underwent a band-pass filter to highlight the cellular nuclear morphology.

We utilized BYOL^18^ for advanced self-supervised image feature extraction. In BYOL, the images were resized to 224 × 224 pixels and subjected to two random augmentations: horizontal flipping, color distortion, Gaussian blurring, and solarization. These images were fed into two networks: the online network (comprising an encoder, projector, and predictor) and the target network (similar but without the predictor). Both networks used the ResNet^53^ architecture for encoding and multilayer perceptrons for projecting and predicting. Embeddings from these networks were synchronized using an *L*2 loss for consistency.

After training, the encoder of the target network produced a 2,048-dimensional feature vector for each image, which was reduced to 64 components using Principal Component Analysis^54^. The resulting matrix *C* ∈ ℝ^*N*×64,^ represented the image features, with each row *c*_*i*_ corresponding to the features of spot *v*_*i*_.

### Integration of slices

Each slice was processed individually, while dealing with multiple consecutive tissue slices. Given *m* slices with the respective numbers of spots *N*_1_, *N*_2_, …, *N*_*m*,_ the total spot count *T* is the sum of all *N*_*i*_. We then obtained distinct adjacency matrices *A*^1^ to *A*^*m*^ and image feature matrices *C*^1^ to *C*^*m*^ for each slice. Batch integration commenced with the vertical concatenation of raw gene expression into a single matrix, which was then normalized to a mean of zero and scaled for unit variance. From this, *F* HVGs were selected, forming the integrated expression matrix *χ* ∈ ℝ^*T*×*F*^.

In the next step, we integrate these matrices. The adjacency matrices were combined into a block-diagonal matrix *𝒜*, as shown in Equation (1), and the image feature matrices were vertically concatenated to form matrix *𝒞*, as shown in Equation (2): These integrated matrices formed the basis for further analysis.

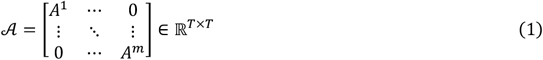

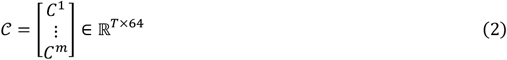

### Graph augmentation

The contrastive learning workflow in the STAIG is based on the GCA framework^55^. Each iteration augmented the input graph, resulting in two modified graphs, *G*^1^ and *G*^2^, and their associated gene expression matrices, *𝒳*^1^ and *𝒳*^2^. These were derived from the original graph *G*, which possessed an edge set *ℰ*, through a specific augmentation strategy applied to the integrated adjacency matrix *𝒜* and gene expression matrix *𝒳*.

Utilizing the integrated image feature matrix *𝒞*, we computed the image spatial distance matrix *D*^*img*^ ∈ *R*^*T*×*T*^, with 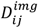 indicating the Euclidean distance between spots *v*_*i*_ and *v*_*j*_ in the image feature space. The edge weight matrix *W* is defined by combining *𝒜* with *D*^*img*^, followed by a logarithmic transformation to modulate the distance values, as expressed in Equation (3).

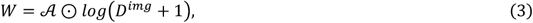

The symbol ⊙ represented the element-wise multiplication, and the addition of 1 prevented the logarithm of zero.

*W* was then scaled to a probability matrix *P* ∈ ℝ^*T*×*T*^ by the SoftMax function, with each element *P*_*i,j*_ being calculated as per Equation (4):

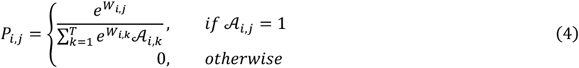

During each iteration, the augmented graphs *G*^1^ and *G*^2^ maintained the node set *V* of original graph *G*. However, their sets of edges, *ℰ*^1^ and *ℰ*^2^ were independently formed by probabilistically removing each edge from *ℰ* (for example, an edge (*v*_*i*_, *v*_*j*_) ∈ *ℰ*), based on the probability 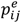, derived from *P* and adjusted within a range set by *p*_*min*_ and *p*_*max*_ as in Equation (5).

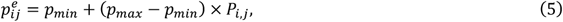

Here, *p*_*min*_ represents the minimum probability that an edge will be considered for removal. The thresholds for *p*_*min*_ and *p*_*max*_ were set according to Eq (6).

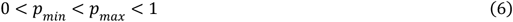

In scenarios lacking image data, STAIG defaults to a fixed edge removal probability of 0.3.

The gene expression matrix *𝒳* was augmented by introducing variabilities such as salt-and-pepper noise into the images. This involves two independent random vectors, *r*^1^ and *r*^2^, each within {0,1}^*F*^. The elements in the vectors are sampled using a Bernoulli distribution. The augmented matrices *𝒳*^1^ and *𝒳*^2^ are then formulated by multiplying each row *x*_*i*_ in *𝒳* by *r*^1^ and *r*^2^ respectively, as described in Equation (7).

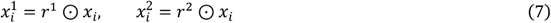

Here, 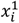and 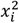corresponded to the respective rows in *𝒳*^1^ and *𝒳*^2^.

### Contrastive learning in the STAIG framework based on the GCA architecture

In every iteration, *G*^1^ and *G*^2^ are processed through a shared GNN structured with *l* layers of GCNs, followed by two layers of fully connected networks, yielding embeddings *H*^1^ and *H*^2^ for each graph as follows:

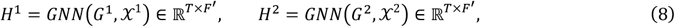

where *F*^′^ ≪ *F*, and the rows 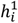 and 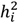 in *H*^1^ and *H*^2^ represented the embeddings of spot *v*_*i*_ in the augmented graphs *G*^1^ and *G*^2^, respectively.

In the STAIG model, neighbor contrastive learning loss^56^ was employed to ensure the consistency and distinctiveness of the embeddings in *H*^1^ and *H*^2^. This involved constructing and comparing positive and negative sample pairs.

For positive pairs, we considered the embedding 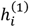of *v*_*i*_ in *G*^1^ as an example. Positive pairs were sourced from: **(1)** The embedding 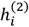of the same spot *v*_*i*_ in *G*^2^; **(2)** embeddings of neighboring nodes of *v*_*i*_ in *G*^1^, denoted as 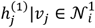, where 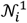 represented the set of neighboring nodes of *v*_*i*_ in *G*^1^; and **(3)** embeddings of neighboring nodes of *v*_*i*_ in *G*^2^, expressed as 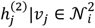, with 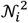being the set of neighbors of *v*_*i*_ in *G*^2^. The total number of positive pairs for anchor 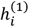is 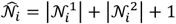, where 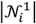and 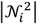denote the counts of neighboring nodes of *v*_*i*_ in *G*_1_ and *G*_2_, respectively.

By contrast, negative sample pairs are generally defined as all pairs other than the positive ones. However, selecting negative samples without considering class information led to a sampling bias^19^, where many negative samples were incorrectly classified as belonging to the same class. This negatively affected the clustering results. To address this issue, when the images were available, we implemented the DS^19^. DS filters out false negatives based on the similarity of the image features *c*_*i*_ of each spot *v*_*i*_. Initially, the spots were clustered into *Q* classes using the k-means method based on their image features, creating pseudo-labels 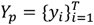. During the construction of negative samples, any *v*_*i*_ sharing the same pseudo-label were excluded.

The neighbor contrastive loss associated with anchor 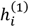 between *G*^1^ and *G*^2^ is given by

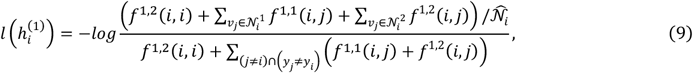

here, 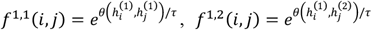, *τ* was a temperature parameter, and *θ*(·) represented a similarity measure (in our work, the inner product is employed). Given the structural similarity between *G*^1^ and *G*^2^, computation of 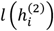mirrored that of 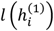. The final neighbor contrastive loss averaged over both augmented graphs is expressed as

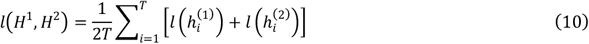

### Spatial domain identification through clustering and refinement

Upon completion of training, the spot embeddings in both augmented graphs were averaged to derive the final low-dimensional representation *H* given as follows:

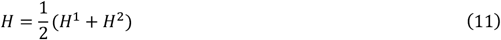

The *H* was clustered using the mclust^57^ algorithm based on a Gaussian Mixture Model. For datasets with manual annotations, the number of clusters was set to match the number of labeled classes. In the absence of such annotations, SC determines the number of clusters, with the count corresponding to the highest SC designated as the final number for the spatial domain.

Consistent with previous baseline methods, we employed a refinement technique for the DLPFC dataset^20^ to further mitigate noise and ensure smoother delineation of clustering boundaries. As part of this refinement, for a given *v*_*i*_, all spots within a radius *d* in its spatial vicinity were assigned to the same class as *v*_*i*_.

Regarding multi-slice integration, because all data were amalgamated into a single matrix prior to model input, the embedding of spots across different slices outputted by the GNN eliminated batch differences. The procedure for spatial identification mirrored that of single-slice processing.

### The overall architecture of STAIG

The architecture of the STAIG features a GNN with one GCN layer and two layers of a fully connected network, producing a 64-dimensional *F′*. It employed a Parametric Rectified Linear Unit activation function and an Adam optimizer with an initial learning rate of 5 × 10^−4,^ and a weight decay of 1 × 10^−5^. The model ran for 400 epochs, with a temperature parameter τ set to 10 and a neighbor count *k* of 5 by default. Further details on hyperparameter optimization can be found in the ‘Hyperparameter optimization’ of the Supplementary Information.

### The ablation studies of STAIG

To assess the effectiveness of the STAIG based solely on its novel contrastive learning framework and the efficacy of the graph augmentation strategy utilizing image information, we conducted ablation experiments on the DLPFC dataset. The experimental setup included a basic framework without image data, using a fixed graph edge removal probability and DS with image data, employing only adaptive random edge removal based on image features, and implementing a complete framework.

The results demonstrated the consistency of STAIG’s superiority in the median ARI compared with other methods, even in the absence of image data. Both the adaptive random edge removal strategy and DS were grounded in significantly improved average and median ARI values. Comprehensive details of these ablation studies are provided in the ‘Ablation studies’ of the Supplementary Information.

### Visualization and functional analysis of spatial domain

The detected spatial domains were visualized using UMAP. Differential gene expression was analyzed using the Wilcoxon test for SCANPY. Gene ontology analysis was done Cluster clusterProfiler^58^. For differentially expressed gene analysis, |log2FC| > 0.25 and p-value < 0.05 were used as the marker gene selection thresholds. P-values were calculated using the Wilcoxon rank-sum test.

## Data availability

The datasets examined in our study can be accessed in their unprocessed states directly from the creators, as detailed in Supplementary Tables S1 and S2. Additionally, the data utilized in this study are now openly accessible at [https://doi.org/10.5281/zenodo.10437391].

## Code availability

The source code for STAIG is openly available for academic and noncommercial use. This information can be accessed from the following GitHub: [https://github.com/y-itao/STAIG]. This repository includes all necessary instructions for installation, usage, and a detailed README for guidance. It is also deposited at Zenodo [https://doi.org/10.5281/zenodo.10437391].

## Declarations

## Acknowledgments

Computational resources were provided by the supercomputer system SHIROKANE at the Human Genome Center, the Institute of Medical Science, the University of Tokyo.

## Funding

This study was supported by the JSPS KAKENHI [22K06189 to K.N., JP22K21301 to M.L., and 20H05940 to S.P.].

## Authors’ contributions

Y. Y. and K. N. conceived and designed the study. Y.Y. and Y.C. performed the data analysis. Y.Y., Y.C., and K.N. drafted the manuscript. S.P., M.L., X.Z., and Y.Z. provided guidance for data analysis. All the authors have read and approved the final version of the manuscript.

## Ethics approval and consent to participate

Not applicable.

## Consent for publication

Not applicable.

## Competing interests

Not applicable.

## Notes

### Competing Interest Statement

The authors have declared no competing interest.

### Summary of Updates

revised phrasing and an updated list of authors

